# *Drosophila melanogaster* employs nuclear architecture for recruitment of dosage compensation

**DOI:** 10.1101/2024.10.29.620935

**Authors:** Maggie P. Lauria Sneideman, Victoria H. Meller

## Abstract

*Drosophila melanogaster* males increase expression from their single X chromosome to match the expression from two female X chromosomes. Increased expression involves recruitment of the Male Specific Lethal (MSL) complex to X-linked genes and modification of chromatin by this complex. How X-linked genes are selectively identified remains unclear, but *cis*-acting recruiting elements are involved. Chromatin Entry Sites (CES) bind an adapter protein that directly recruits the MSL complex. In addition, a class of AT-rich satellites, the 1.688^X^ repeats, are enriched on the X and facilitate compensation of nearby genes. How these elements cooperate to identify the X is unknown. Nuclear architecture influences dosage compensation in many organisms. In this study we test the idea that factors involved in nuclear architecture participate in recruitment by the CES and 1.688^X^ repeats. We find genetic interactions between mutations that reduce X recognition and factors that govern chromatin structure, silencing, remodeling and insulation, suggesting roles in X recognition. A reporter for recruitment revealed that Heterochromatin Protein 2 (HP2) and Scaffold Attachment Factor-A (SAF-A) disrupt compensation but do not selectively influence recruitment by either element. In contrast, Imitation switch (ISWI), D1 and Nucleoporin 153kD (Nup153) contribute to recruitment by the CES and Centrosomal protein 190kD (Cp190) exerts a striking influence on recruitment by 1.688^X^ repeats. These findings reveal that recruitment by the CES and 1.688^X^ repeats involves nuclear organization. We propose that the use of *cis*-acting elements that engage complementary aspects of nuclear organization ensures robust X recognition.

## INTRODUCTION

*Drosophila melanogaster* females have two gene-rich X chromosomes and males have a single X and a gene-poor Y. This creates an imbalance in the ratio of X to autosomal gene products that is lethal if uncorrected (Disteche, 2012). *Drosophila* males increase expression from X-linked genes to match that in the female, a process termed dosage compensation. This is accomplished in part by a complex of proteins and RNA, the Male Specific Lethal (MSL) complex, that is recruited to transcribed X-linked genes (Gelbart and Kuroda, 2009). One protein in the complex, Males absent on the first (MOF), is a histone acetyltransferase that modifies histone H4 on lysine 16 (H4K16ac) (Hilfiker, 1997; Akhtar and Becker 2000). H4K16ac decompacts chromatin to facilitate initiation and elongation (Shogren-Knaak et al., 2006; Conrad et al., 2012). H4K16ac is abundant within the bodies of X-linked genes in males, where it contributes to increased expression (Copur et al., 2016; Ferrari 2013). Loss of any protein in the MSL complex blocks complex activity and is male-lethal (Lucchesi and Kuroda, 2015). The long non-coding RNAs, *RNA on the X 1* and *-2* (*roX1, roX2*) are essential but redundant components of the MSL complex. Loss of both *roX* RNAs leads to mislocalization of the MSL proteins, reduced expression of X-linked genes and male lethality (Meller and Rattner, 2002; Deng et al., 2005).

How the MSL complex selectively recognizes the X chromosome is poorly understood. The intact complex is believed to initially recognize about a hundred X-linked Chromatin Entry Sites, also known as High Affinity Sites (CES, HAS) (Kageyama et al., 2001; Kelley et al., 1999; Alekseyenko et al., 2008; Straub et al., 2008). It then spreads to nearby active genes through recognition of the co-transcriptional histone H3 lysine 36 trimethylation mark by the Male Specific Lethal 3 (MSL3) subunit, ensuring enrichment of the MSL complex in active genes (Larschan et al., 2007; Sural et al., 2008; Bell et al., 2008; Buscaino et al., 2006; Kind and Akhtar, 2007; Alekseyenko et al., 2006; Gilfillan et al., 2006). A breakthrough came with the identification of Chromatin Linked Adapter Protein (CLAMP), a pioneer transcription factor that binds GA-rich MSL recognition elements (MREs) within the CES and recruits the MSL complex (Soruco et al., 2013; Alekseyenko et al., 2008). CLAMP also fulfills essential functions in both sexes, including formation of the Histone Locus Body on the second chromosome (Rieder et al., 2017).

Autosomal sites of CLAMP binding fail to recruit the MSL complex, indicating that additional factors guide X recognition. Some CES have been found to contain an extension of the binding site that interacts with MSL2 (Eggers et al., 2023). Termed PionX sites, these achieve earlier recruitment of the complex and are enriched on the X chromosome (Villa et al., 2016). Although the different classes of MRE-containing CES are central elements in X recognition, additional factors also contribute to selective recognition of X-linked genes.

We previously demonstrated that AT-rich satellite repeats enriched in X euchromatin facilitate recruitment of the MSL complex to nearby genes (Joshi and Meller, 2017; Deshpande and Meller, 2018). These repeats are ∼359 bp in length with a buoyant density of 1.688 gm CsCl/cm^3^ and are collectively referred to as the 1.688^X^ repeats (DiBartolomeis et al., 1992; Waring and Pollack, 1987; Kuhn et al., 2012). Ectopic expression of small interfering RNA (siRNA) from 1.688^X^ repeats enhances the survival of *roX1 roX2* males and increases X localization of MSL proteins (Menon et al., 2014; Biswas et al., 2024). Autosomal insertions of 1.688^X^ DNA enable nearby, active genes to recruit the MSL complex by a mechanism involving the siRNA pathway (Joshi and Meller, 2017). Recruitment is enhanced by 1.688^X^ siRNA (Deshpande and Meller, 2018). Although CLAMP is necessary for recruitment by the CES, it is unnecessary for recruitment by 1.688^X^ repeats (Makki and Meller, 2024). Likewise, the siRNA pathway is unnecessary for recruitment by the CES. We have proposed that the CES and 1.688^X^ satellites act in complementary ways to ensure selective X recognition.

Dosage compensation occurs within the context of a highly organized nucleus. Spatial proximity and nuclear organization are factors in identification of compensated chromatin in flies and other organisms (Ramirez et al., 2015; Sharma et al., 2014; Lau et al., 2014; Splinter et al., 2011; Chaumeil et al., 2006; Chen et al., 2016; Teller et al., 2011; Cerase et al., 2019). In support of this idea, the CES engage in long range interactions that are proposed to help define the X territory (Grimaud and Becker, 2009; Ramirez et al., 2015). It is unclear whether the 1.688^X^ repeats rely on nuclear organization to recruit compensation, but 1.688^X^ repeats have high potential as Matrix/Scaffold Attachment Regions (MAR/SAR) and participate in long range interactions suggesting this possibility (Pathak et al., 2014; Stroud et al., 2020). The role of nuclear organization in recruitment by the 1.688^X^ satellites, and how different facets of nuclear organization influence CES and 1.688^X^ function, remain unexplored.

We selected candidate genes that bind satellite DNA, remodel chromatin or participate in other aspects of nuclear organization. As dosage compensation is essential in males, we reasoned that knockdown of some factors involved in this process might produce male-biased lethality, but this was not observed in otherwise wild type flies. In contrast, several knockdowns did enhance the male-lethality of partial loss of function *roX1 roX2* chromosomes. The signature defect in *roX1 roX2* males is disrupted X-localization of the MSL complex, suggesting that these factors might contribute to X recognition. An assay for recruitment of compensation revealed that that two of the genes identified act non-specifically to disrupt the process of dosage compensation, but others selectively influence recruitment by the CES or 1.688^X^ repeats. Our findings highlight the role of nuclear organization in recruitment. We propose that CES and 1.688^X^ repeats engage complementary aspects of nuclear organization to achieve selective X recognition.

## METHODS

### Fly strains

Ribonucleic acid interference (RNAi) knockdown strains and mutations are presented in Supplemental Table 1. *roX1 roX2* mutations were previously described (Deng et al., 2005; Menon and Meller, 2014). The P{*sqh*-GAL4} driver was a gift from Dr. S. Todi (Wayne State University). We acknowledge FlyBase (Öztürk-Çolak et al., 2024) and the Bloomington *Drosophila* Stock Center (NIH P40OD018537) for providing essential resources and *Drosophila* strains.

### Expression measurements

Knockdown efficiency was determined by qRT-PCR of cDNA from third instar larvae as described by Koya and Meller (2015) (see Supplemental Table 2 for primers). In brief, groups of 50 larvae were homogenized in Trizol and RNA extracted as recommended by the manufacturer (Invitrogen). RNA was cleaned using the QIAGEN RNeasy Kit. One μg cleaned RNA was reverse transcribed with random hexamers and SuperScript IV (Invitrogen). Expression was measured by qRT-PCR of technical duplicates for three biological replicates. Knockdown efficiency was calculated by the ΔΔCt method with normalization to DCTN2-p50 (*dmn*) (Pfaffl, 2001). A Student’s t-test determined significance. To measure the effect of D1 knockdown on repeat expression, D1 knockdown line P{TRiP.JF03031} was mated to the P{*sqh*-GAL4} driver and larvae collected. Primers that amplify 1.688^1A^, 1.688^3F^, and the 359 repeats were designed to amplify repeats at cytological positions 1A, 3F and closely related repeats that comprise 11 Mb of X heterochromatin (359 repeats) (Supplemental Table 2). Template was diluted 1:160 with water to measure the high multiplicity 359 bp repeats. Repeat expression was normalized to *dmn* and compared to a *yw* laboratory reference strain.

### Knock down and detection of genetic interactions

To determine the effect of knock down in otherwise wild type flies, females heterozygous for each RNAi transgene were mated to homozygous driver males (P{*sqh*-GAL4}). RNAi offspring were distinguished by the *y^+^* marker and survival was calculated by the ratio of *y^+^* to *y* flies. Matings were done in triplicate and a Student’s t-test was used to determine significance.

To detect genetic interactions, *yw roX1^ex33^ roX2Δ*; P{*sqh*-GAL4} virgins were mated to *yw* males heterozygous for knockdown transgenes. Knockdown adults were identified by body color and survival calculated as described above. Matings, performed in triplicate, produced a total of 500-1000 offspring for each RNAi line. A Student’s t-test was used to determine significance.

### Reporter for recruitment of compensation

An autosomal reporter for recruitment of compensation was created by placing 1.688^X^ repeats, *roX1* with an internal CES (*roX1*/CES) or both recruiting elements in a transgene containing Firefly luciferase (Fluc) (Makki and Meller, 2024). Males containing these Fluc reporters, a *Renilla* luciferase normalizer and P{*sqh*-GAL4} were mated to knockdown virgins. Groups of three one day old adult males were frozen at –20°C for at least 30 min and crushed in 300 μl lysis buffer (1% Triton X-100, 10% glycerol, 25 mM glycylglycine pH 7.8, 15 mM MgSO_4_, 4 mM EGTA, 1 mM DTT). Lysates were briefly centrifuged to remove debris and 50 μl aliquots frozen at –20°C.

Twenty μl lysate was analyzed in a 96-well plate microplate (F-bottom, Greiner) using a GloMax microplate luminometer (Promega). One hundred μl of Firefly luciferase reagent (25 mM glycylglycine (Fisher, cat # AC120140250), 15 mM K_X_PO_4_ (mixture of monobasic dihydrogen phosphate and dibasic monohydrogen phosphate, pH 8.0), 4 mM EGTA, 2 mM ATP, 1 mM DTT, 15 mM MgSO_4_, 0.1 mM Coenzyme A (Nanolight, #309), 75 μM luciferin (Nanolight, #306), pH 8.0) was dispensed and luminescence read for ten seconds after a two second delay. One hundred μl of Renilla luciferase buffer (1.1 M NaCl, 2.2 mM EDTA, 0.22 M K_X_PO_4_ (pH 5.1), 0.44 mg/mL BSA, 2 mM NaN_3_, 10 μM coelenterazine (Nanolight, #303), pH adjusted to 5.0) was added and emission read for ten seconds following a two second delay. Activity was recorded as relative light units (RLU) and emission with lysis buffer alone was subtracted from all values. Values were normalized by dividing Firefly RLU by Renilla RLU. Normalized values for Firefly luciferase in the absence of recruiting elements or knock down was set to 1.

### Polytene preparation and immunostaining

Salivary glands from wandering third instar male larvae were dissected in Ringer’s solution, fixed in PBS with 4% formaldehyde and 1% Triton-X 100 for 45 seconds and moved to 50% acetic acid, 4% formaldehyde for 2 minutes. Glands were moved to 50% acetic acid, 16.66% lactic acid, 4% formaldehyde on a cover slip and squashed. Preparations were frozen in liquid nitrogen, cover slips removed and stored at –20°C in 95% ethanol until use.

Preparations were rehydrated for 30 minutes in PBST (1X PBS, 0.2% Triton-X 100) and blocked in PBST with 0.2% BSA for 30 minutes. Thirty μl of a 1:200 dilution of anti-MSL-1 serum in block (gift of M. Kuroda, Harvard University) was placed on each preparation, covered with parafilm and incubated overnight at 4°C in a humidified chamber. Slides were washed three times with PBST. Forty μl of preabsorbed Texas Red conjugated to anti-rabbit in blocking solution (1:200 dilution) was placed on each slide, covered with parafilm and incubated in a dark, humidified chamber for 3 hours. Slides were washed three times with PBST, mounted with ProLong Diamond Antifade Mountant (Life Technologies Corporation) and images photographed at 100X (Leica Microsystems, DM5500 B). DAPI and Texas Red channels were overlaid using ImageJ.

## RESULTS

### Selection and evaluation of candidate genes

We chose a candidate gene approach to test the idea that nuclear organization participates in recruitment of dosage compensation by CES and 1.688^X^ repeats (Supplemental Table 1). Reasoning that AT-hook proteins might directly bind the AT-rich 1.688^X^ repeats, we selected HP2, an AT-hook and heterochromatin-associated protein that interacts with HP1 (Schaffer et al., 2006). HP2 also interacts with the ISWI-containing Nucleosome Remodeling Factor (NURF), which regulates *roX* transcription and is recruited to CES (Bai et al., 2007; Tsukiyama et al., 1995). Another AT-hook protein, D1, localizes to 359 bp repeats that make up ∼11 Mb of pericentric X heterochromatin and are closely related to the euchromatic 1.688^X^ repeats (Alfageme et al., 1980; Aravind and Landsman, 1998; Blattes et al., 2006; Sproul et al., 2020). We examined factors involved in chromatin structure, including Scaffold Attachment Factor-A (SAF-A; also known as heterogeneous nuclear ribonucleoprotein U, hnRNPU) and Scaffold Attachment Factor-B (SAF-B). These factors are broadly involved in maintenance of chromatin structure, and each possesses features suggesting a possible role in recruitment by the CES or 1.688^X^ repeats. Both proteins contain a SAP domain, proposed to be involved in DNA binding (Gohring et al., 1997). SAF-A also has an RNA binding domain and binds both RNA and DNA in mammalian cells (Nozawa et al., 2017). *Drosophila* SAF-B interacts with specific sites on polytene chromosomes and is a component of the insoluble nuclear matrix (Alfonso-Parra & Maggert, 2010). Wild type SAF-B localization depends on the SAP domain, transcriptional activity and RNA.

Subunits of the nuclear pore and nuclear envelope were also examined. Loss of the nuclear pore proteins Megator (Mgtor) and Nucleoporin 153kD (Nup153) disrupt localization of the dosage compensation complex and Mgtor knockdown increases X-linked gene expression, suggesting a role in governing the level of activation (Mendjan et al., 2006; Aleman et al., 2021). The nuclear envelope protein Lamin B influences chromatin architecture and interacts with Ago2, a protein that localizes to some 1.688^X^ repeats and participates in recruitment of compensation (Nazer et al., 2018; Menon and Meller, 2012; Deshpande and Meller, 2018; Makki and Meller, 2024). We also selected the Lamin B receptor (LBR) as it anchors chromatin to the nuclear envelope (Wagner et al., 2003; Bondarenko and Sharakhov, 2020).

Finally, we examined the representative insulator proteins Modifier of mdg4 (Mod(mdg4)), an enhancer of variegation that interacts with CLAMP, and Centrosomal protein 190kD (Cp190), a zinc finger protein found at most insulators and some boundaries (Bag et al., 2019; Gerasimova et al., 2007; Gerasimova et al., 1995; Pai et al., 2004; Whitfield et al., 1995).

Knockdown efficiency under the strong, constitutive P{*sqh*-GAL4} driver was determined (Franke et al., 2005; Supplemental Fig. 2). Knockdown lines that did not achieve detectable reduction in target mRNA, or were fully lethal, were not pursued further (Supplemental Table 1). As dosage compensation is an essential, male-limited process we induced knockdown in otherwise wild type flies but found no significant differences between male and female survival (Fig. 1). Full survival was observed with most knock downs, but a 90% reduction in SAF-A mRNA reduced recovery of male and female adults by about 50%, and a 47% reduction in Cp190 mRNA reduced adult survival by 40%. Notably, we failed to observe reduced male survival upon ISWI knockdown. ISWI is essential in both sexes, but some ISWI mutations reduce male survival and disrupt the morphology of the polytenized male X chromosome (Deuring et al., 2000; Corona et al., 2002). The 58% mRNA reduction we achieved permits full male survival (Supplemental Fig. 2).

**Figure 1.**
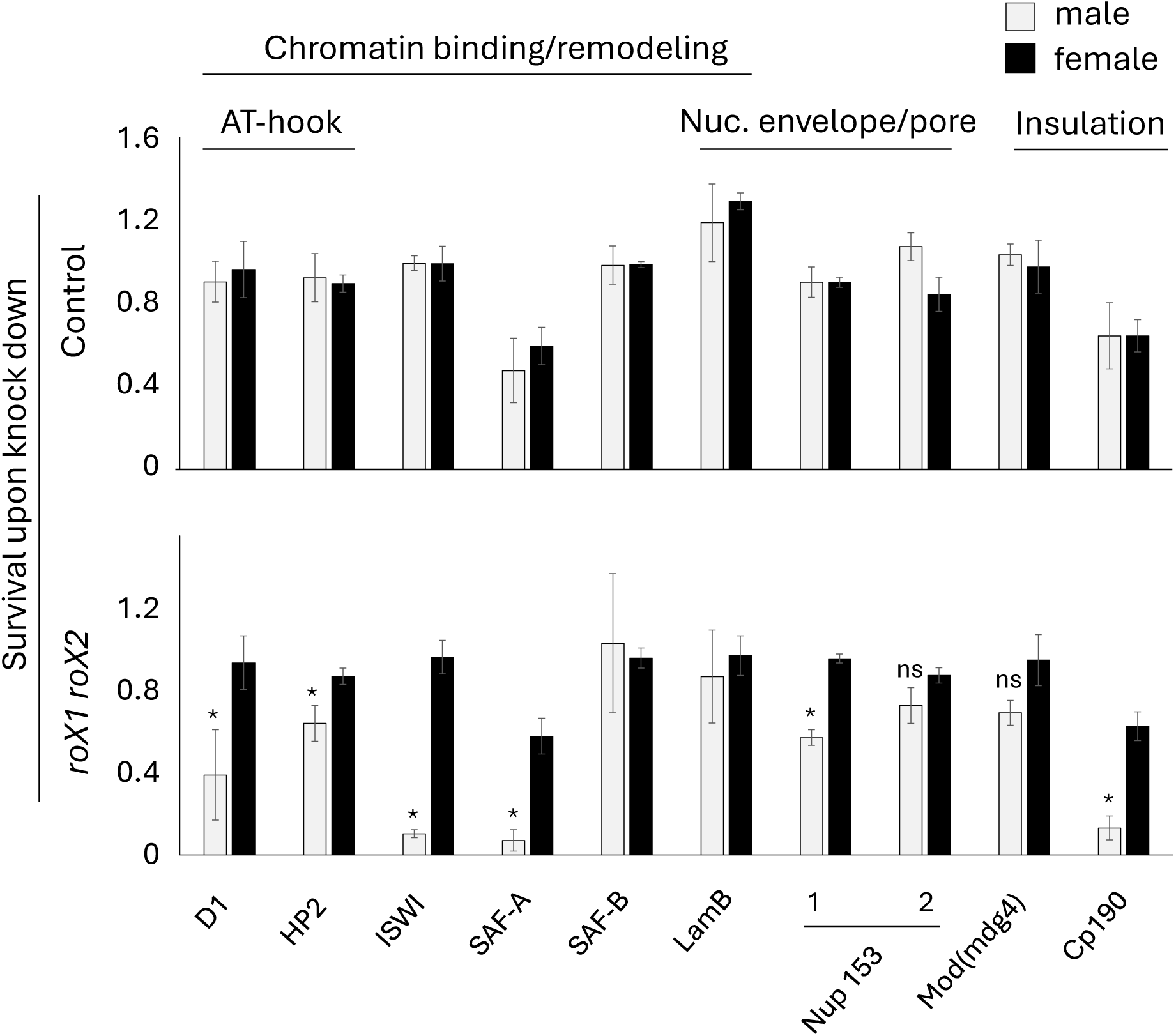
Factors that mediate nuclear organization enhance *roX1 roX2* male lethality. (Top) Knockdown of factors involved in nuclear organization does not produce male-biased lethality. Survival is calculated by dividing the number of knockdown offspring by control offspring. (Bottom) Knockdown in *roX1^ex33^ roX2Δ* males identifies genes that may play a role in X recognition. Survival is calculated by dividing knockdown adults by siblings of the same sex without knockdown. The average of three replicates is presented. Significance was determined by comparing knockdown flies to siblings without knockdown using a Student’s t-test. Error bars signify standard error. * p-value < 0.05. (D1 P{TRiP.JF03031}; HP2 P{TRiP.JF01304}; ISWI P{TRiP.JF01582}; SAF-A P{TRiP.HMC03927}; SAF-B P{TRiP.HMC05536}; LamB P{TRiP.JF01389}; Nup153 1 P{TRiP.HM05248}; Nup153 2 P{TRiP.HMS00527}; Mod(mdg4) P{TRiP.HMS00849}; Cp190 P{TRiP.HMJ02105}).

At least one *roX* RNA is necessary for normal MSL localization to the X chromosome (Meller and Rattner, 2002). Because the signature defect in *roX1 roX2* males is failure of X-recognition, the *roX1^ex33^ roX2Δ* X chromosome, allowing ∼30% adult male escapers, is a sensitized background to identify genes involved in this process. We mated *roX1^ex33^ roX2Δ*; P{*sqh*-Gal4} females to males heterozygous for knockdown transgenes marked with *y^+^*. All sons are *roX1^ex33^ roX2Δ,* carry the P{*sqh*-Gal4} driver and those undergoing knockdown are identified by *y^+^* body color. Female survival reflects the effect of knockdown alone as females are heterozygous for *roX1^ex33^ roX2Δ* and do not dosage compensate.

Knockdown of D1, ISWI, SAF-A, and Cp190 produced striking reductions in *roX1 roX2* male survival, and knockdown of Nup153 and HP2 produced modest but significant reductions in male survival (Fig. 1). ISWI serves as a proof of concept as the NURF complex has an established role at the CES (Bai et al., 2007). The remainder of our study focuses on the six factors that permit survival to adulthood and show genetic interactions with *roX1 roX2*.

### ISWI knockdown reduces recruitment by the CES

We utilized a luciferase reporter to determine if genes displaying genetic interactions with *roX1 roX2* contribute to recruitment by the 1.688^X^ repeats or CES (Makki and Meller, 2024). Recruiting elements, either *roX1* with an internal CES (*roX1*/CES), 1.688^X^ repeats or both elements, are placed next to Firefly luciferase (Fluc) and inserted on an autosome. Fluc activity with no recruiting element reflects basal activity (Fig. 2a). Fluc expression is normalized to *Renilla* luciferase (Rluc) situated on a different autosome. Flies homozygous for Fluc transgenes, P{*sqh*-GAL4} and Rluc were mated to laboratory reference (*yw*) flies and luciferase activity measured in adult male offspring (Fig. 2a-d). The Fluc/Rluc ratio increased 4-5 fold when recruiting elements were present (Fig. 2g). This exceeds the two-fold increase anticipated for full compensation, as has been previously observed at *roX1*/CES transgenes (Kelley and Kuroda, 2003). Hyperactivation is thought to stem from disruption of normally repressive chromatin by recruitment of the MSL complex. Knockdown of a factor necessary for recruitment by *roX1*/CES, or the 1.688^X^ repeats, prevents activation by the respective recruiting element (Fig. 2e,f). To determine if engaging the RNAi pathway itself influences luciferase activity, we knocked down genes with no connection to MSL recruitment in each reporter strain. Knockdown of *yellow* (*y*), with an endogenous target, or mCherry, which induces futile RNAi, failed to affect reporter activity (Fig. 2g; see also Makki and Meller, 2024).

**Figure 2.**
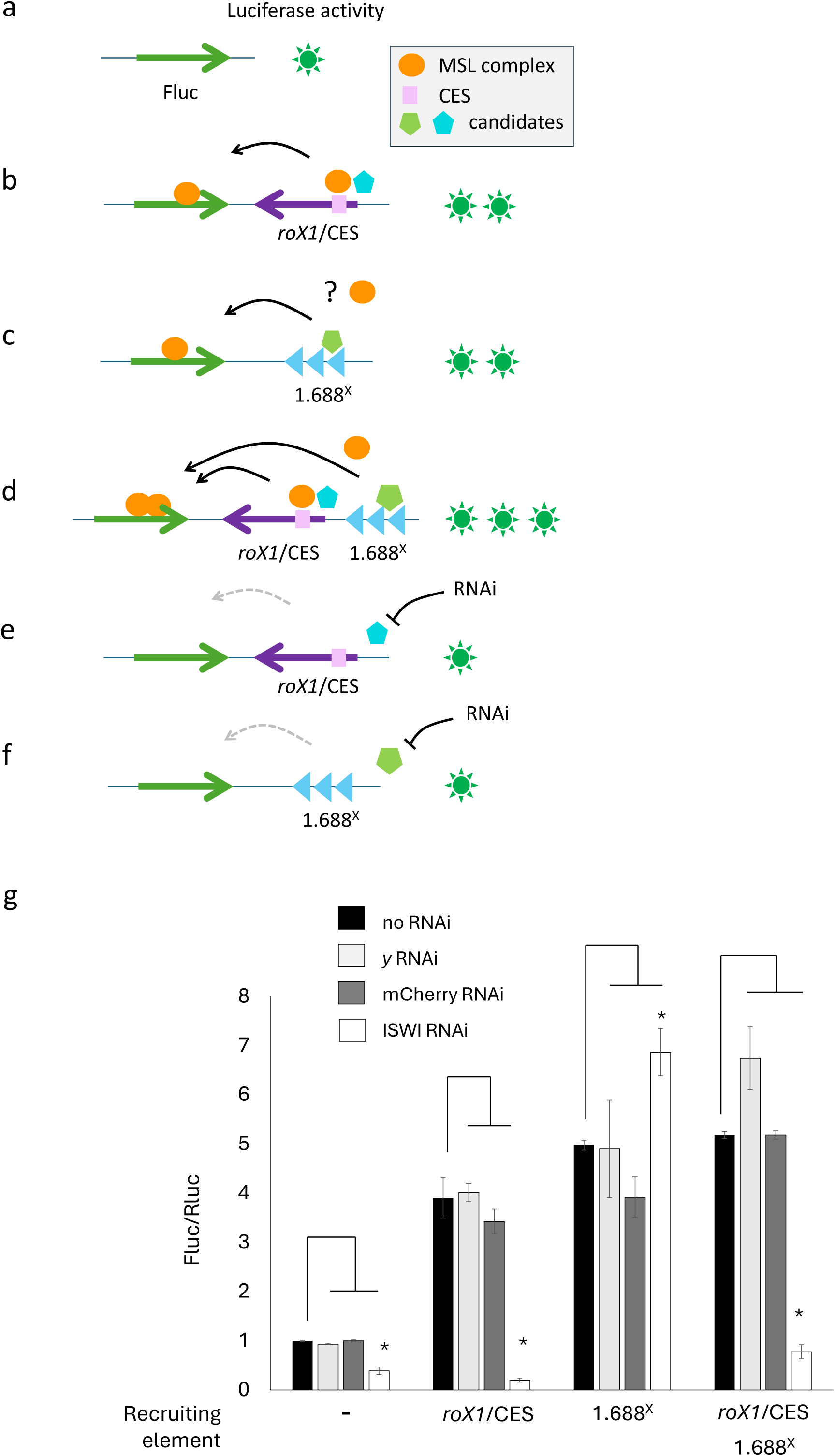
ISWI knockdown decreases recruitment by *roX1*/CES. (a) Constitutive Firefly luciferase (Fluc) is driven by the *dmn* promoter (not shown). A low level of Fluc activity (green star) is detected. b) The MSL complex (orange) is recruited by factors that interact with *roX1*/CES (turquoise). The MSL complex spreads into Fluc and increases expression. c) 1.688^X^ repeats recruit the MSL complex to nearby active genes through a mechanism involving distinct factors (green). d) Transgenes with *roX1*/CES and 1.688^X^ repeats achieve the highest activation of Fluc. e) Knockdown of a protein that acts through the CES will prevent activation when *roX1*/CES is the recruiting element. f) Knockdown of a factor acting through 1.688^X^ repeats will block activation. g) Control knockdown does not influence Fluc activity, but ISWI knockdown blocks recruitment only by *roX1*/CES. The Fluc/Rluc ratio with no recruiting element or knockdown is set to 1. The significance of three biological replicates was determined by a Student’s t-test. * = p<0.05. Error bars represent standard error. (ISWI P{TRiP.JF01582}; *y* P{TRiP.HMC05546}; mCherry P{VALIUM20-mCherry.RNAi}).

We then examined luciferase activity in males knocked down for ISWI. In agreement with a role for the ISWI-containing NURF complex at the CES, knockdown produced a striking decrease in Fluc activation by the *roX/*CES alone, or when both recruiting elements were present (Fig. 2g; Bai et al., 2007). In contrast, there was a modest increase in activity when 1.688^X^ repeats were the sole recruiting element. A reduction in Fluc activity was observed when no recruiting elements were present, an effect that may reflect the genome-wide role of ISWI in regulation of chromatin accessibility (Deuring et al., 2000). We conclude that ISWI conforms to our expectations for a general chromatin factor that also serves a specific role in recruitment of compensation by the CES.

### Disruption of chromatin structure sensitizes males to a defect in dosage compensation

Knockdown of HP2 achieved modest but significant increases in the expression of firefly luciferase regardless of the presence or nature of recruiting elements (Fig. 3). This is consistent with the idea that HP2 fulfills a repressive function but fails to identify a specific role for HP2 in recruitment by the *roX1*/CES or 1.688^X^ repeats. Conversely, SAF-A knockdown dramatically decreased firefly luciferase expression from all Fluc reporters, consistent with the proposed role of SAF-A in maintenance of an open chromatin structure that facilitates expression (Fig. 3) (Nozawa et al., 2017). Although knockdown of SAF-A or HP2 did not selectively influence recruitment by either roX1/CES or 1.688^X^ repeats, knockdown did lower *roX1 roX2* male survival. This is consistent with the idea that disruption of chromatin compromises the function of the MSL complex in a manner that does not selectively influence recruitment through *roX1*/CES or the 1.688^X^ repeats.

**Figure 3.**
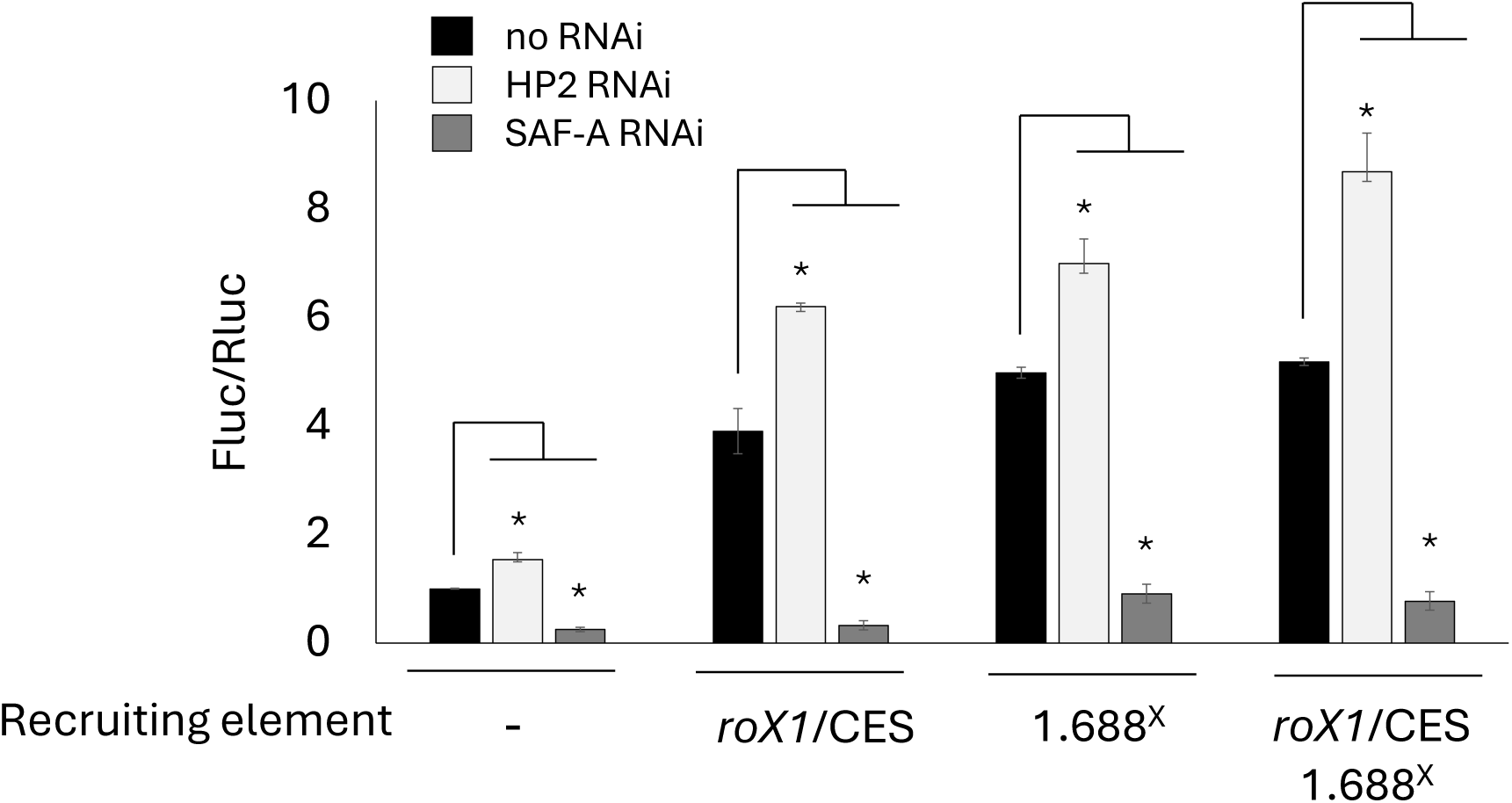
SAF-A and HP2 modulate expression in a general manner. HP2 knockdown (light gray) increases Fluc expression regardless of the presence or nature of recruiting elements. In contrast, SAF-A knockdown (dark gray) sharply decreases Fluc expression. Bars represent standard error of three biological replicates. Significance was calculated with the Student’s t-test (*p<0.05). (HP2 P{TRiP.JF01304}; SAF-A P{TRiP.HMC03927}).

### D1 promotes recruitment through roX1/CES

The association of D1 with AT-rich heterochromatic 359 bp satellites suggested that D1 might also bind the closely related euchromatic 1.688^X^ repeats. Unexpectedly, knockdown of D1 achieved a striking decrease in recruitment by the *roX1/*CES but not by 1.688^X^ repeats (Fig. 4). As Fluc activation is unaffected when 1.688^X^ repeats are the sole recruiting element, D1 knockdown does not block the ability of the MSL complex to enter and activate chromatin at active genes. It is possible that D1 acts directly at the CES, but more likely that knockdown of D1 may produces indirect effects by altering chromatin over the 359 bp repeats, comprising 11 Mb of pericentric heterochromatin. Reduction or displacement of D1 reduces PEV of some variegating insertions, indicating a reduction in silencing (Aulner et al., 2002; Janssen et al., 2000). However, D1 knockdown failed to appreciably influence accumulation of transcripts from heterochromatic 359 bp repeats or from 1.688^X^ clusters at 1A and 3F (1.688^1A^, 1.688^3F^; Supplemental Fig. 3). While the influence of D1 knockdown on *roX1*/CES recruitment is striking, the underlying mechanism remains speculative.

**Figure 4.**
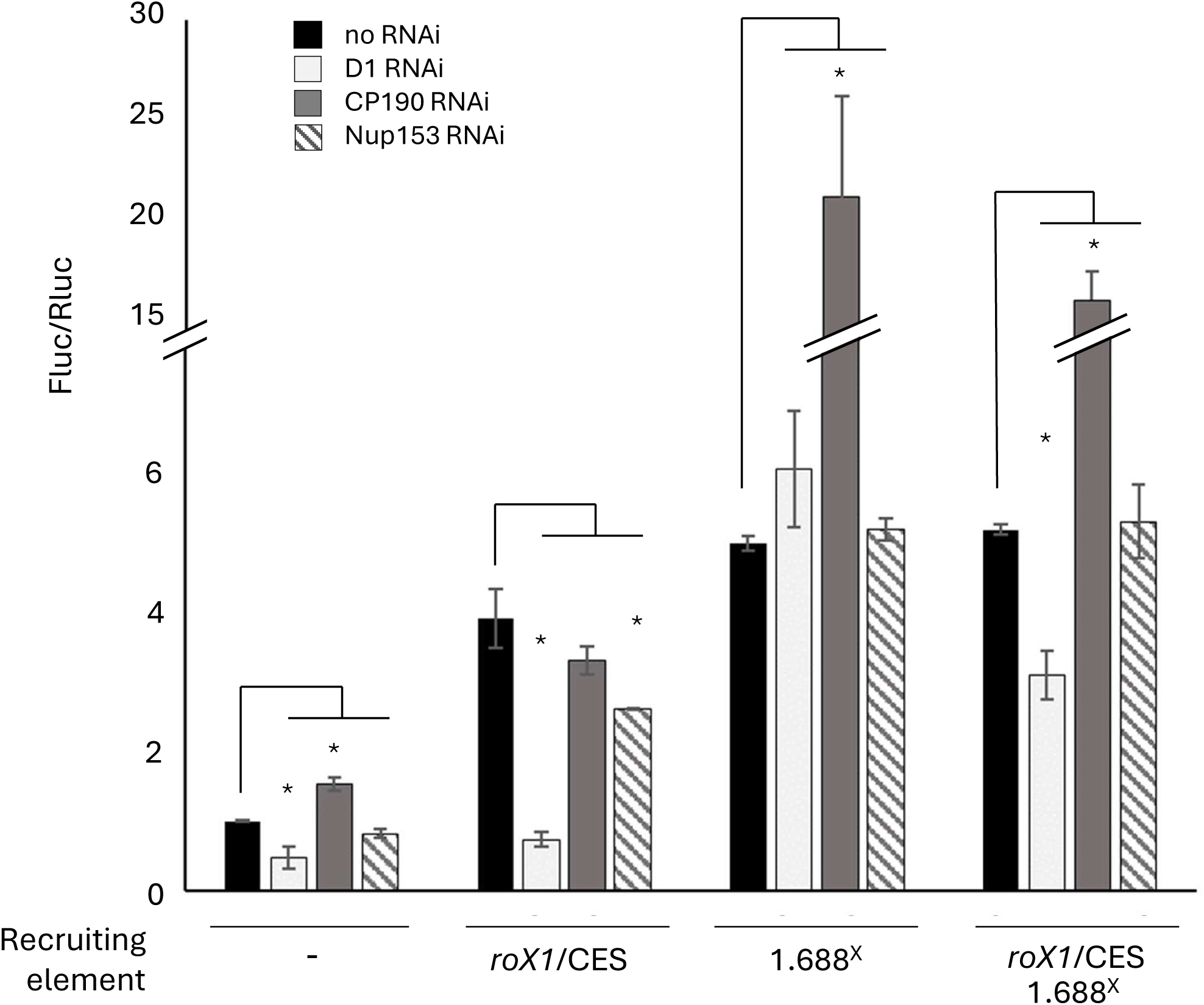
D1, Cp190, and Nup153 influence individual recruiting elements. D1 knockdown (light gray) reduced Fluc activity without recruiting elements and when *roX1*/CES was present. Cp190 knockdown (dark gray) increased Fluc expression slightly with no recruiting element but achieved a dramatic increase when 1.688^X^ repeats were present. Nup153 knockdown (hatched) decreased Fluc activity when *roX1*/CES was present. Error bars represent standard error of three biological replicates. Significance was calculated with the Student’s t-test (*p<0.05). (D1 P{TRiP.JF03031}; Cp190 P{TRiP.HMJ02105}; Nup153 P{TRiP.HM05248}).

### Nup153 modulates recruitment by the roX1/CES

MOF has been found to associate with the nuclear pore protein Nup153, and a 70-80% reduction of Nup153 in tissue culture cells blocked X-localization of the MSL complex, leading the conclusion that Nup153 facilitates MSL binding to the X chromosome (Mendjan et al., 2006). Consistent with this, we found that a 50% reduction in Nup153 mRNA achieved a modest reduction in Fluc expression by the *roX1*/CES recruiting element (Fig. 4). In contrast, knock down had no effect on recruitment through 1.688^X^ repeats, or when both recruiting elements were present. Although the interaction of Nup153 and MOF suggests a direct recruitment, MOF also participates in the Non-Specific Lethal (NSL) complex, a regulatory complex that includes some of the MSL proteins and is found at many housekeeping genes (Lam et al., 2012). Although it is possible that the MOF-Nup153 interaction occurs in the context of a different complex, our current study reveals that Nup153 contributes to recruiting by the CES.

### Cp190 limits recruitment by 1.688^X^ repeats

Cp190 is known to interact with CLAMP, leading us to predict a role in recruitment through the *roX1*/CES (Bag et al., 2019). Surprisingly, Cp190 knockdown had no influence on Fluc expression when *roX1*/CES was the sole recruiting element but induced a 15 to 20-fold increase in Fluc expression when 1.688^X^ was present alone, or with *roX1*/CES (Fig. 4). Cp190 is recruited by DNA-binding factors that play central roles in nuclear organization, often in the context of insulation or the establishment of borders (Mohan et al., 2007; Kaushal et al., 2022; Kahn et al., 2023; Ramirez et al., 2018; Bag et al., 2021). However, activation of Fluc upon Cp190 knockdown is only observed when 1.688^X^ repeats are present, linking the function of these repeats to a factor that mediates insulator activity and long range interactions.

### Immunodetection on polytene chromosomes reveals disruption of recruitment

We next turned to polytene preparations to evaluate the effect of D1, Cp190 and Nup153 knock down on MSL1 localization. Knockdown was done in otherwise wild type or in *roX1^ex40^ roX2Δ* males. *roX1^ex40^* retains sufficient *roX1* sequence to support high male survival but has an internal deletion of 2.3 kb (Deng et al., 2005). Although MSL1 localization to the X remains strong in *roX1^ex40^ roX2Δ* males, ectopic location at autosomal sites, including the chromocenter and telomeres, is elevated. We have found *roX1^ex40^ roX2Δ* to be a sensitive background for evaluation of factors that influence MSL localization to polytene chromosomes (Menon and Meller, 2014). Knock down of D1 alone, or with *roX1^ex40^ roX2Δ,* had little discernable influence on localization to the X chromosome (Fig. 5a,b). This is surprising, given that D1 knock down reduces the survival of *roX1^ex33^ roX2Δ* males and dramatically reduces Fluc expression when the *roX1*/CES recruiting element is present (Fig. 1).

**Figure 5.**
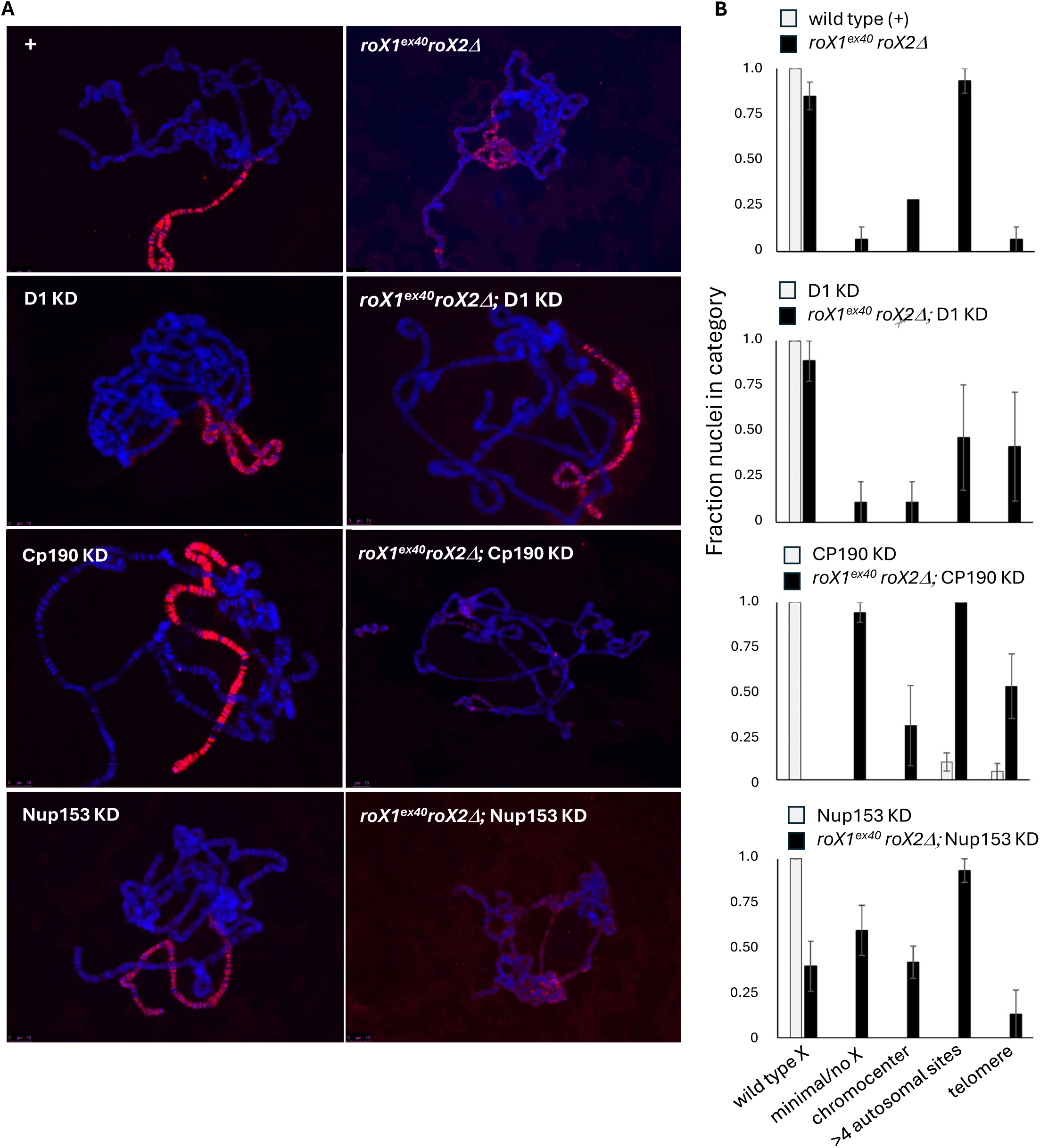
MSL1 immunodetection reveals disruption of X recognition. a) Representative images of polytene preparations from male larvae. Left panels are control or knockdown alone, right panels are knockdown in *roX1^ex40^ roX2D* males. MSL1 is detected by Texas Red and DNA by DAPI (blue). (b) Polytene preparations were scored for X localization indistinguishable from wild type (wild type X); minimal or no X localization and ectopic MSL1 localization to the chromocenter (chromocenter); four or more ectopic autosomal MSL1 sites (>4 autosomal sites); or telomeres (telomere). Eleven to 22 nuclei were scored on 3 slides of each genotype with labels obscured. Error bars represent standard error. (D1 P{TRiP.JF03031}; Cp190 P{TRiP.HMJ02105}; Nup153 P{TRiP.HM05248}).

Although X localization of MSL1 remains strong in Cp190 knockdown males, a slight increase in autosomal localization was observed (Fig. 5b; compare wild type and Cp190 knock down). In contrast, no wild type MSL1 binding is observed in *roX1^ex40^ roX2Δ* males upon Cp190 knockdown. Ectopic MSL1 binding to autosomal sites is prominent in all preparations and few adult males emerge from these matings, indicative of near-complete synthetic lethality. Taken together, these findings support the idea that Cp190 normally participates in localization of MSL complex to the X chromosome.

While Nup153 knockdown alone did not appreciably disrupt the pattern of MSL1 localization or increase ectopic binding, the intensity of signal over the X chromosome appeared subjectively lower in many preparations (Fig. 5a). When knockdown of Nup153 was done in *roX1^ex40^ roX2Δ* males, the fraction of nuclei with a wild type pattern of X localization decreased and those with minimal or no X localization sharply increased from that observed in *roX1^ex40^ roX2Δ* (Fig. 5). These observations are consistent with a role for Nup153 in recruitment of the MSL complex to the X chromosome, as has been previously reported (Mendjan et al., 2006; Vaquerizas et al., 2010).

## DISCUSSION

Dosage compensation is a model epigenetic system, but our understanding of how organisms selectively identify a single chromosome is incomplete. We find that factors that regulate nuclear architecture act in complex ways to support compensation of the fly X chromosome. The candidates examined here can be divided into three classes based on the response of recruiting elements to knock down (Fig. 6). Knockdown of the first group, comprised of HP2 and SAF-A, reduced the survival of *roX1 roX2* males but did not selectively influence recruitment by *roX1*/CES or the 1.688^X^ repeats. SAF-A knockdown reduced Fluc expression regardless of the presence or nature of recruiting element. SAF-A is thought to maintain interphase chromatin structure by linking DNA with non-coding and nascent RNAs to form an open meshwork that permits transcription (Nozawa et al., 2017). Consistent with this model, mutation of SAF-A or RNase treatment in mammalian cells condenses chromatin (Hall et al., 2014). These effects are most apparent in expressed, gene-rich regions that display chromatin decondensation, a cytological feature of active genomic regions (Creamer et al., 2021). The single male X is partially decondensed, appearing the same width as the two paired female X chromosomes in polytene preparations (Dobzhansky 1957). It is possible that SAF-A contributes to this structural difference. Interestingly, mammalian SAF-A anchors *Xist*, a long non-coding RNA that coats the inactive female X chromosome to induce dosage compensation (Hasegawa *et al.,* 2010; Kolpa *et al.,* 2016; Helbig and Fakelmayer, 2003).

**Figure 6.**
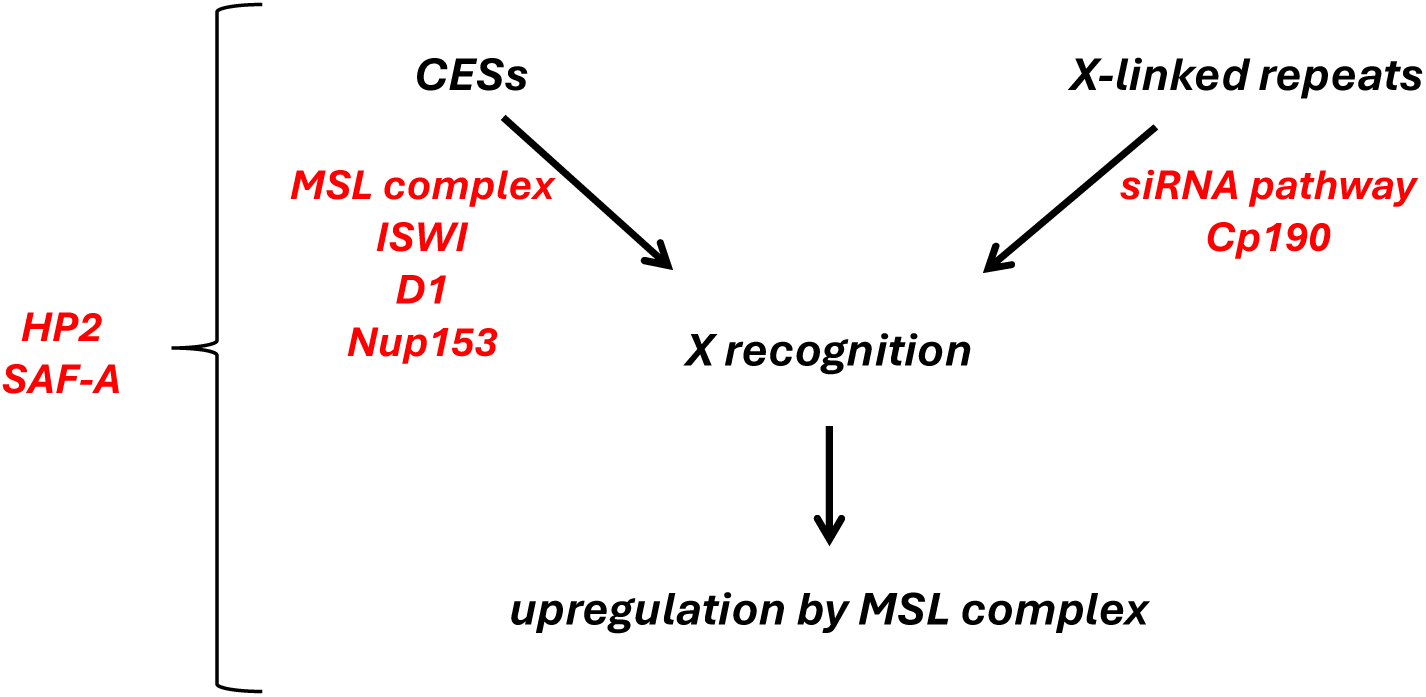
Contribution of architectural factors to dosage compensation. Genes of interest examined for recruiting activity. The CES in *roX1* binds CLAMP to directly recruit the MSL complex. Activities of ISWI, D1 and Nup153 support recruitment by the CES. Recruitment by 1.688^X^ repeats requires the siRNA pathway and is inhibited by Cp190, although this may be indirect. HP2 and SAF-A have general effects on chromatin organization. We propose that males with a defect in dosage compensation have heightened sensitivity to loss of these factors.

HP2 attracted our attention because of AT-hooks suggesting the potential for association with AT-rich 1.688^X^ DNA. HP2 associates with heterochromatic proteins and is enriched in heterochromatin at telomeres and the centromere (Mendez et al., 2013; Alekseyenko et al., 2014; Shaffer et al., 2006, Stephens et al., 2006). Mutations in some genes necessary for heterochromatin formation produce disordered banding of the polytenized male X chromosome (Spierer et al., 2005; Deuring et al., 2000). A minor enrichment of HP1 on the X chromosome also suggests a link between heterochromatic factors and the process of dosage compensation (De Wit et al., 2005; Liu et al., 2005; Park et al., 2019). The finding that knockdown of HP2 increased Fluc expression regardless of recruiting elements is consistent with a general repressive role for HP2, but the genetic interaction with *roX1 roX2* suggested a function in dosage compensation. We postulate that reduction of SAF-A or HP2 sensitizes males to defects in dosage compensation. One way this might occur is if the MSL complex is unable to function correctly in an altered chromatin environment.

We also identified factors that manage chromatin state, structure or insulation and contribute to recruitment by *cis*-acting elements more selectively. Knock down of ISWI, D1 or Nup153 reduced recruitment by the *roX1*/CES but had little effect on recruitment by 1.688^X^ repeats. ISWI, previously linked to dosage compensation and CES function, serves as a proof of concept (Corona et al., 2002; Bai et al., 2007). Previous studies revealed that ISWI fulfills multiple functions related to dosage compensation. In partial loss of function ISWI mutants the polytenized X chromosome becomes short, bloated and the banding pattern is lost, but MSL complexes continue to localize to the X and acetylate H4 (Badenhorst et al., 2002; Corona et al., 2002). Normal polytene morphology is restored by mutations that block H4K16ac deposition, suggesting that ISWI restores chromatin compaction following activation by MSL complex activity (Corona et al., 2002). In addition, the ISWI-containing NURF complex has been found to localize to the CES and support spreading of the MSL complex from the CES (Bai et al., 2007). An autosomal *roX2* transgene in a NURF mutant background accumulates MSL complex around the insertion site, producing an expanded region of heavy MSL localization. Our study confirms that ISWI participates in recruitment of dosage compensation by the *roX1*/CES and distinguishes the mechanism of recruitment by *roX1*/CES from that of the 1.688^X^ repeats, which continue to recruit upon ISWI knock down.

In this study we also identified additional factors that act at *roX1*/CES, including Nup153. The genetic interaction between Nup153 and *roX1 roX2* validates previous work identifying a role for Nup153 in dosage compensation (Mendjan et al., 2006; Vaquerizas et al., 2010). MSL localization was blocked by a 70-80% reduction of Nup153 protein in *Drosophila* SL-2 cells (Mendjan et al., 2006). In contrast, the 43% reduction of Nup153 mRNA we achieved in flies is benign in otherwise wild type males but, when combined with *roX1^ex40^ roX2Δ,* exacerbates MSL1 mislocalization. Both Mgtor and Nup153 colocalize to dosage compensated genes, are pulled down by association with MOF and necessary for localization of the MSL complex to the X chromosome (Mendjan et al., 2006; Vaquerizas et. al., 2010). Interestingly, a previous study found that Mgtor played a role in limiting the level of X activation (Aleman et al., 2021). Indeed, Mgtor knock down partially rescued *roX1 roX2* males. Transcriptional activation is frequently associated with recruitment to the nuclear pore (reviewed in Gasser and Ahktar, 2007). However, many pore proteins are not exclusive to the nuclear envelope but found in association with chromatin (Reviewed in Capelson, 2023). The finding that Nup153 acts at *roX1*/CES implicates Nup153, and potentially nuclear pore localization, in CES function.

Knock down of D1 also blocked recruitment by *roX1*/CES but not the 1.688^X^ repeats. D1 is a multi-AT-hook protein that localizes to pericentric 359 bp repeats found on the X chromosome (Blattes et al., 2006). As the heterochromatic 359 bp and euchromatic 1.688^X^ repeats are closely related, we hypothesized that D1 might also contribute to the function of the 1.688^X^ repeats (Sproul et al., 2020). It was therefore unexpected that D1 knock down selectively blocked recruitment by *roX1*/CES. Mutation of D1 disrupts the chromocenter and reduces nuclear envelope integrity, suggesting a general breakdown of nuclear organization (Jagannathan et al., 2019). It is possible that recruitment by *roX1*/CES is particularly sensitive to the defect in nuclear organization produced by loss of D1. One way this might occur is if nuclear disorganization impairs long range interactions between CES are enhance recruiting activity. It is also possible that loss of D1 produces a shift in the availability of other proteins recruited to 359 bp repeats, and that this exerts an indirect effect on *roX1*/CES recruiting. Despite uncertainties about the mechanism responsible, the finding that D1 influences recruitment by *roX1*/CES explains the genetic interaction between D1 and *roX1 roX2*.

The current study identified one factor, Cp190, with a powerful influence on recruitment through the 1.688^X^ repeats. The specificity of Cp190 for 1.688^X^ repeats was surprising in light of studies finding that Cp190 and CLAMP, which binds the CES in *roX1*, are mutually dependent for chromatin binding (Bag et al., 2019). Polytene chromosomes revealed a striking reduction in MSL1 localization to the X chromosome when knockdown was performed in the *roX1^ex40r^oX2Δ* background. This accounts for genetic interactions between Cp190 and roX mutants but fails to address the mechanism involved. Cp190 is recruited by a number of DNA binding factors to mediate enhancer-promoter interactions (Kaushal et al., 2022; Chathoth et al., 2022; Liang et al., 2014). Cp190 also participates in the long-range interactions that contribute to insulation and boundary formation (Vogelmann et al., 2014; Mourad and Cuvier, 2016; Hou et al., 2012). Insulation serves to limit enhancer action to the intended target, and loss of Cp190 produces modest increases in expression when enhancers access nearby but previously inaccessible promoters (Schwartz et al., 2012). A modest activation of Fluc is observed in the absence of recruiting elements, consistent with this idea. But no increase is observed when *roX1*/CES is the sole recruiting element, indicating that the dramatic activation of Fluc upon Cp190 knockdown is related to the specific function of the 1.688^X^ repeats. The 1.688^X^ repeats are depleted for transcription factors but subject to siRNA-directed H3K9me2 deposition and enriched for Ago2 (Roy et al., 2010; Deshpande and Meller, 2018). Interestingly, Ago2 interacts with Cp190 in some insulators (Moshkovich et al., 2011). It is possible that Cp190 limits the recruiting activity of 1.688^X^ through a direct interaction with elements of the siRNA pathway. If so, this would provide a remarkable opportunity to dissect the mechanism of recruitment by 1.688^X^ repeats, which is at present speculative. It is also possible that depletion of Cp190 releases factors that act through 1.688^X^ repeats but are in limiting supply. The list of Cp190 interactors contains many intriguing candidates, but the magnitude of Fluc activation upon a relatively modest, 50% knock down of Cp190 argues against this possibility.

In summary, we have shown that males with defect in X recognition have heightened sensitivity to disruption of chromatin state and nuclear architecture. We find that factors involved in chromatin structure, remodeling, DNA binding and insulation influence specific recruiting pathways, sometimes in unexpected ways (Fig. 6). These findings reveal a complex relationship between recruiting elements and nuclear organization. The fact that some architectural factors selectively influence 1.688^X^ or CES recruiting suggests that normal identification of X chromatin involves collaboration between different aspects of nuclear organization.

### Limitations of this study

Reliance on a single Fluc reporter insertion site raises the possibility of confounding effects due to local chromatin elements. However, this reporter has faithfully recapitulated the roles of numerous factors that were known or suspected to act at *roX1*/CES or 1.688^X^ repeats. We chose to limit analysis to non-lethal knockdown strains that enabled evaluation of genetic interactions, X chromosome histology and reporter expression. Adapting the luciferase reporter system to accommodate knockdown strains of varying efficiencies is a future goal.

## ACKNOWLEDGEMENTS

We thank the VanBerkum lab at Wayne State for the use of their fluorescent microscope. We also thank Dr. Reem Makki for development of the luciferase assay and assistance with it. Stocks obtained from the Bloomington *Drosophila* Stock Center (NIH P40OD018537) were used in this study. This work was supported by a Wayne State University summer dissertation award to MPLS and NIH award ROGM093110 to VHM.

