## supplemental figures for "*Drosophila melanogaster* employs nuclear architecture for recruitment of dosage compensation"

### Supplemental Material

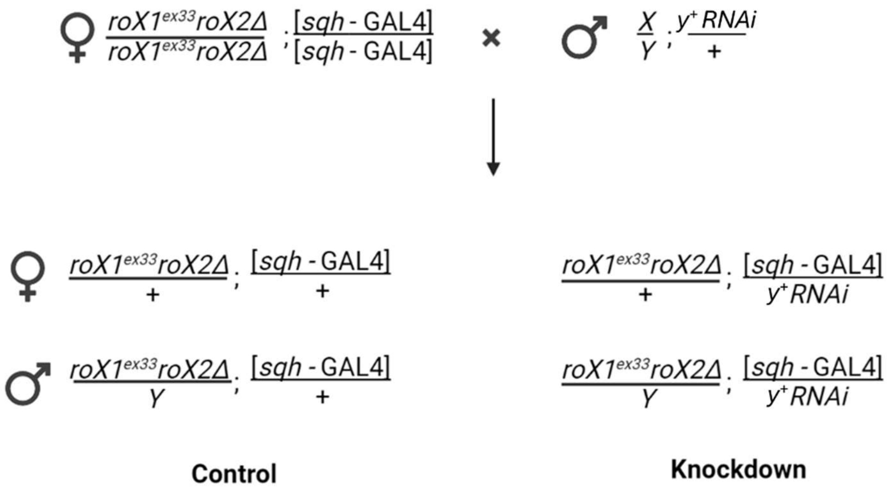


**Supplemental Figure 1. Mating to test candidate gene knockdown in *roX1 roX2* males**. Abundance of control and knockdown offspring is compared to determine a genetic interaction.

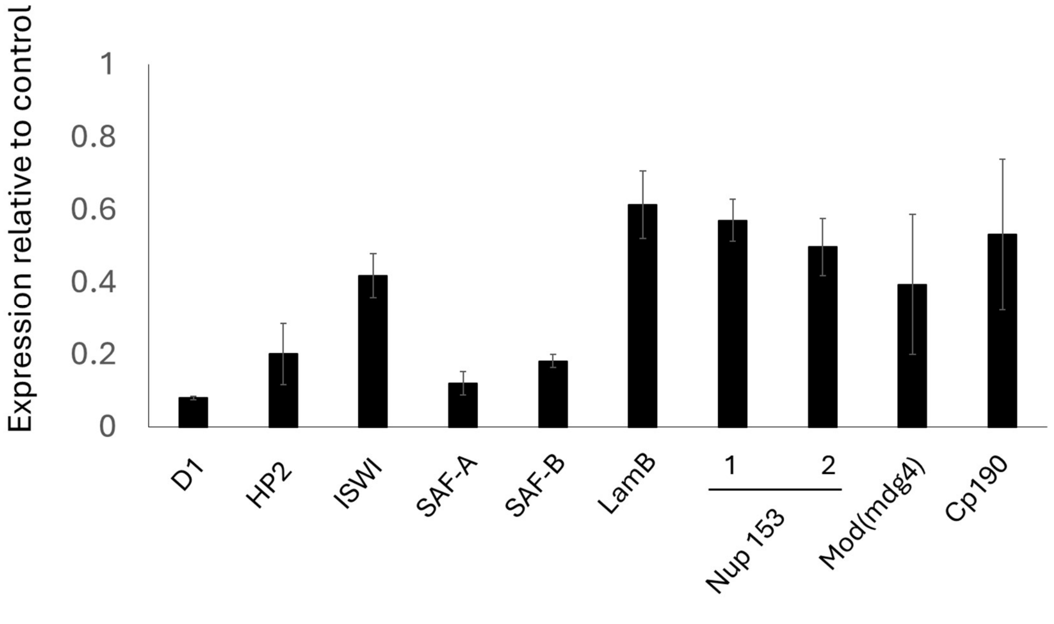


**Supplemental Figure 2. Estimation of knockdown efficiency by qPCR.** Error bars represent standard error of three biological replicates. Expression levels relative to control *yw* expression levels. (D1 P{TRiP.JF03031}; HP2 P{TRiP.JF01304}; ISWI P{TRiP.JF01582}; SAF-A P{TRiP.HMC03927}; SAF-B P{TRiP.HMC05536}; LamB P{TRiP.JF01389}; Nup153 1 P{TRiP.HM05248}; Nup153 2 P{TRiP.HMS00527}; Mod(mdg4) P{TRiP.HMS00849}; Cp190 P{TRiP.HMJ02105}).

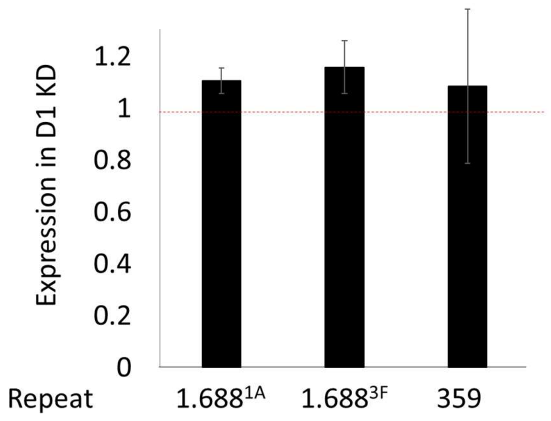


**Supplemental Figure 3. Knockdown of D1 does not alter expression from several repeats.** (RNAi line D1 P{TRiP.JF03031}). Primers listed in Supplemental Table 2. Error bars represent standard error of three biological replicates.

**Supplemental Table 1. Fly strains used in these studies**. "roX" indicates a genetic interaction with *roX1 roX2*. "Leth." Indicates this line is lethal upon RNAi knockdown. "Inc." indicates inconclusive result.

| Protein | Stock # | FBID | Genotype | % KD | roX | Leth. | Inc. |
|---|---|---|---|---|---|---|---|
| Cp190 | 42536 | FBst0042536 | $y^1$ $v^1$; P{TRiP.HMJ02105}attP40 | 46.8 | x | | |
| Cp190 | 33903 | FBst0033903 | $y^1$ $sc^*$ $v^1$ $sev^{21}$; P{TRiP.HMS00845}attP2 | - | | x | |
| D1 | 28616 | FBgn0000412 | $y^1$ $v^1$; P{TRiP.JF03031}attP2 | 91.9 | x | | |
| D1 | 33655 | FBgn0000412 | $y^1$ $sc^*$ $v^1$ $sev^{21}$; P{TRiP.HMS00061}attP2 | 26 | | x | |
| D1 | 17340 | FBst1029357 | P{VT019579-lexA::GADfl}attP40 | - | | | x |
| D19a | 33371 | FBgn0022935 | $y^1$ $sc^*$ $v^1$ $sev^{21}$; P{TRiP.HMS00244}attP2 | 48.3 | | | x |
| HDAC1 | 34846 | FBst0034847 | $y^1$ $sc^*$ $v^1$ $sev^{21}$; P{TRiP.HMS00165}attP2 | - | | | x |
| HP2 | 31346 | FBgn0026427 | $y^1$ $v^1$; P{TRiP.JF01304}attP2 | 79.7 | x | | |
| HP2 | 38255 | FBgn0026427 | $y^1$ $sc^*$ $v^1$ $sev^{21}$; P{TRiP.HMS01699}attP40 | 74.5 | | x | |
| ISWI | 31111 | FBgn0011604 | $y^1$ $v^1$; P{TRiP.JF01582}attP2 | 58.1 | x | | |
| ISWI | 51931 | FBgn0011604 | $y^1$ $v^1$; P{TRiP.GLC01788}attP40 | 27.8 | | | x |
| LamB | 31605 | FBgn0002525 | $y^1$ $v^1$; P{TRiP.JF01389}attP2 | 38.6 | | | x |
| LBR | 63269 | FBgn0034657 | P{TRiP.HMC02426}attP2 | 88.1 | | | x |
| LBR | 43417 | FBst0043417 | $w^*$; P{GSV1}$LBR^{EP-439}$ | - | | | x |
| Mgtor | 32941 | FBgn0034657 | $y^1$ $sc^*$ $v^1$ $sev^{21}$; P{TRiP.HMS00735}attP2 | 67.5 | | x | |
| Mgtor | 10537 | FBst0010537 | $y^1$ $w^{67c23}$; P{lacW}$Mgtor^{k03905}$/CyO | - | | | x |
| mod(mdg4) | 33907 | FBgn0002781 | $y^1$ $sc^*$ $v^1$ $sev^{21}$; P{TRiP.HMS00849}attP2 | 60.6 | | | x |
| mod(mdg4) | 32995 | FBgn0002781 | $y^1$ $sc^*$ $v^1$ $sev^{21}$; P{TRiP.HMS00795}attP2 | 51.2 | | x | |
| Nup153 1 | 30504 | FBgn0061200 | $y^1$ $sc^*$ $v^1$ $sev^{21}$; P{TRiP.HM05248}attP2 | 42.9 | x | | |
| Nup153 2 | 32837 | FBgn0061200 | $y^1$ $sc^*$ $v^1$ $sev^{21}$; P{TRiP.HMS00527}attP2 | 50.2 | | | x |
| Nup98-96 | 28562 | FBst0028562 | $y^1$ $v^1$; P{TRiP.HM05048}attP2 | - | | x | |
| SAF-A | 55209 | FBgn0050122 | $y^1$ $sc^*$ $v^1$ $sev^{21}$; P{TRiP.HMC03927}attP40 | 87.9 | x | | |
| SAF-B | 64518 | FBgn0039229 | $y^1$ $sc^*$ $v^1$ $sev^{21}$; P{TRiP.HMC05536}attP40 | 81.7 | | | x |

**Supplemental Table 2. Table of primers used for qPCR**

| Target gene | F/R | Sequence |
|---|---|---|
| D1 | F | ACGTTCTTCGATGCTGCTGT |
| | R | CGCCTCTCTATTGTCATCTCG |
| D19a | F | GCCCATTGGCGGTAGTGAAA |
| | R | GATGACTCCGAAGACGGCAC |
| ISWI | F | ATGGGACAGATTGAGCGTGG |
| | R | CTCGCAGCTCTTCGTAGACA |
| SAF-A | F | TTCTTGACCTCGCTCTTGGC |
| | R | AAGCGAATTGCGGTCGTTTG |
| Mod(mdg4) | F | TCGTTATCCGTTAGCCCCTTG |
| | R | TCCAATTCATGTACTGCGGC |
| HP2 | F | GGCTCATCCTCTTCATCGGAG |
| | R | CGCGAACGGTATGAGTCGTT |
| SAF-B | F | TCCGGTGATAAGAAGGACTC |
| | R | CAGTGGAGGATGTGGCAC |
| Mgtor | F | CCAGTTCTCCGCAAAAACAG |
| | R | TTGGCTAGTGCCCACATTTG |
| LamB | F | CCACGGTCAAGAGATAGACC |
| | R | GATTTCCAAATCCAGGGAGAC |
| LBR | F | ATCAAGGCCGCCATCAAG |
| | R | GTTAAATGCTTTGTTCGGTGC |
| Nup153 | F | GAAGCTGCCGAGGGAAG |
| | R | GAAGGCTGGATAGGCTGTTG |
| Cp190 | F | GACAAGAGTACGCCACTAATCAC |
| | R | GGCCCAGTAGTATTCTCTGC |
| 359 rpts | F | TTGTCTGAATATGGAATGTCATATCTC |
| | R | TTCGTTATAACTTGGCTAAAAATGG |
| $1.688^{3F}$ | F | TATTTACAAACGGGGTTATCTCTATAAGG |
| | R | AAAACAGTCTTCATTTAAGCGGTAA |
| $1.688^{1A}$ | F | ATTTACAAACGGGGTTTTCTCTATAACCTG |
| | R | CGTAACAAAATTCCCCATCGACCTG |
